# Brain-state invariant thalamo-cortical coordination revealed by non-linear encoders

**DOI:** 10.1101/148643

**Authors:** Guillaume Viejo, Thomas Cortier, Adrien Peyrache

**Affiliations:** Montreal Neurological Institute, McGill University; École Normale Supérieure, Paris

## Abstract

Understanding how neurons cooperate to integrate sensory inputs and guide behavior is a fundamental problem in neuroscience. A large body of methods have been developed to study neuronal firing at the single cell and population levels, generally seeking interpretability as well as predictivity. However, these methods are usually confronted with the lack of ground-truth necessary to validate the approach. Here, using neuronal data from the head-direction (HD) system, we present evidence demonstrating how gradient boosted trees, a non-linear and supervised Machine Learning tool, can learn the relationship between behavioral parameters and neuronal responses with high accuracy by optimizing the information rate. Interestingly, and unlike other classes of Machine Learning methods, the intrinsic structure of the trees can be interpreted in relation to behavior (e.g. to recover the tuning curves) or to study how neurons cooperate with their peers in the network. We show how the method, unlike linear analysis, reveals that the coordination in thalamo-cortical circuits is qualitatively the same during wakefulness and sleep, indicating a brain-state independent feed-forward circuit. Machine Learning tools thus open new avenues for benchmarking model-based characterization of spike trains.

**Author summary:** The thalamus is a brain structure that relays sensory information to the cortex and mediates cortico-cortical interaction. Unraveling the dialogue between the thalamus and the cortex is thus a central question in neuroscience, with direct implications on our understanding of how the brain operates at the macro scale and of the neuronal basis of brain disorders that possibly result from impaired thalamo-cortical networks, such as absent epilepsy and schizophrenia. Methods that are classically used to study the coordination between neuronal populations are usually sensitive to the ongoing global dynamics of the networks, in particular desynchronized (wakefulness and REM sleep) and synchronized (non-REM sleep) states. They thus fail to capture the underlying temporal coordination. By analyzing recordings of thalamic and cortical neuronal populations of the HD system in freely moving mice during exploration and sleep, we show how a general non-linear encoder captures a brain-state independent temporal coordination where the thalamic neurons leading their cortical targets by 20-50ms in all brain states. This study thus demonstrates how methods that do not assume any models of neuronal activity may be used to reveal important aspects of neuronal dynamics and coordination between brain regions.

## Introduction

Investigating how the brain operates at the neuronal level is usually addressed by the specification of neuronal responses to an experimentally measurable variable or by the quantification of the temporal coordination of neuronal ensembles [Harris, 2005, Rieke, 1999]. Using various methods, the responses of single neurons can be characterized by the tuning curves based on a single measurement (i.e. average firing rate as a function of the observed value) [Hubel and Wiesel, 1962, O’keefe and Nadel, 1978, Taube et al., 1990], with generalized linear models accounting for the coding of multiple features [Harris et al., 2003, Truccolo et al., 2005], biophysical models of spike train generation [Pillow et al., 2005] or information measures and reverse reconstruction [Borst and Theunissen, 1999, Rieke, 1999].

The coding of information in the brain relies on the coordinated firing of neuronal population [Buzsáki, 2010, Harris, 2005, Pouget et al., 2013, Yuste, 2015]. The development of dense electrode arrays [Buzsáki, 2004, Jun et al., 2017] and imaging techniques [Chen et al., 2013, Dombeck et al., 2007] in awake animals now allows monitoring of the activity of large ensembles of neurons and to address fundamental questions about neuronal network coordination. Neuronal interactions, in relation to behavior or internal parameters (e.g. brain states), are evaluated by the statistical dependencies of spike trains, the most widely used method being linear cross-correlations [Perkel et al., 1967]. These linear measures can be generalized to population correlation with tools such as Principal Component Analysis (PCA) [Chapin and Nicolelis, 1999, Peyrache et al., 2010] and Independent Component Analysis [Lopes-dos Santos et al., 2013]. Generalized linear models were used to build predictions of single spike trains as a function of the peer network activity [Harris et al., 2003] and to provide a full statistical description of spatio-temporal neuronal responses and correlations [Pillow et al., 2008]. Methods from graph theory offer ways to compare interactions at the network level across experimental conditions [Humphries, 2017]. Finally, among the large body of available tools, evaluating neuronal coupling by fitting spiking activity to Ising models has provided key insights into the nature of neuronal coordination in a population [Cocco et al., 2009, Schneidman et al., 2006].

The majority of the methods enumerated above rely on a set of assumptions regarding the statistics of the data or the biophysics of neuronal spiking, among others, while seeking explanatory power. To assert the validity of a particular approach, the usual procedure is to divide the data set into a *training set*, used to fit the model parameters, and a *test set*, on which the likelihood of the model is evaluated. However, this method, called cross-validation, does not rule out the possibility that a particular fit of the model parameters, even when leading to high likelihood, corresponds to the wrong model. For example, the omission of a key feature in the model may attribute erroneous contribution to the set of chosen variables. These limitations arise from the lack of ground-truth data that in the most complex (and, therefore, interesting) cases represent an unreachable goal.

This lack of ground-truth data when performing data analysis is particularly unavoidable in neuroscience [Harris et al., 2016]. It has thus become necessary to establish standard, model-free methods that, even if they do not contribute to our understanding of the data, set levels of performance that may be used to benchmark model-based approaches [Benjamin et al., 2017, Truccolo and Donoghue, 2007]. Machine Learning provides a large array of techniques to classify datasets that have demonstrated high level of performance in fields ranging from image processing to astrophysics [LeCun et al., 2015]. Using a supervised classifier, so-called gradient boosting [Benjamin et al., 2017, Truccolo and Donoghue, 2007], we show how this method can determine an encoding model for predicting population spike trains knowing the stimulus input. We also show, in line with recently published work [Benjamin et al., 2017], how gradient boosted trees (XGB) can also be used as a very efficient decoding model that is retrieving the stimulus likelihood knowing the spiking activity of a population of neurons. Finally, we demonstrate how it generates a very accurate encoding model for predicting a population spike train conditioned on another, anatomically projected, set of neuronal activity [Harris et al., 2003].

We tested the validity of the approach on data from the head-direction (HD) system [Peyrache et al., 2015, Taube, 2007, Taube et al., 1990], a sensory pathway whose member neurons, the so-called HD *cells*, emit spike trains that can be explained with high accuracy simply by the direction of the head of the animal in the horizontal plane. Decision trees maximized their branching in input ranges where Fisher Information was maximal. We then determined the optimal parameters of the method for our data set. Finally, we applied this method to simultaneously recorded neurons in the thalamo-cortical network of the HD system, namely in the antero-dorsal nucleus of the thalamus (ADn) and the Post-subiculum (PoSub). We demonstrate that non-linear encoders such as boosted gradients, but not linear analysis, reveal that thalamic neurons lead cortical neurons in a brain-state independent manner.

## Methods

### Gradient boosted trees

Machine Learning literature defines boosting as the combination of many weak classifiers with limited prediction performances in order to build a stronger classifier. The first boosting algorithm is AdaBoost (Adaptive Boosting) [Freund and Schapire, 1995] which trains weak learners using a distribution of weight over the training set. This distribution of weight is updated after the convergence of a weak learner in order for the next weak learner to focus on the difficult examples i.e. the points that are hard to classify.

Boosting algorithms come in different flavors for the type of learners or the updating of the weights [Ferreira and Figueiredo, 2012, Schapire, 2003]. Here we focused on the boosting using the decision tree model as the weak learner. The goal of the gradient boosted trees algorithm is to determine the optimal successive partition of features space in order to assign a weight or a label to a subset of the training examples. This algorithm is thus equivalent to decision trees in which input features are optimally segmented to determine a desired output. The problem is now to apply this reasoning to predict the spiking of neurons based on behavioral features and, conversely, to decode behavioral feature from a population of neurons coding for an internal representation of this feature. Lastly, this algorithms can be useful to predict the spike train of a given neuron from the spiking activity of an upstream neuronal population.

Practically, we first defined the training set [(*x*_1_, *y*_1_),…, (*x*_*m*_, *y*_*m*_)] where *x*_*i*_ ∈ *R*^*d*^ is the i-th training example with *d* different features and *y*_*i*_ is the target value. In this study, we focus on two different types of features: (1) behavioral features, in particular the HD and position of the animals and (2) spiking activity of neuronal ensembles. The goal of the learner reduces to how to make an accurate prediction ŷ_*i*_ given *x*_*i*_ and the correct value *y*_*i*_. A target value *y*_*i*_ for a given training example *x*_*i*_ is a spike count over a finite time bin for one neuron. Assuming neuronal spiking follows an inhomogeneous Poisson distribution, we thus defined the prediction of the model as:

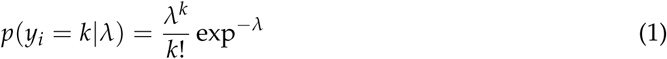

for a given intensity parameter *λ* = *λ*(*x*_*i*_), the single parameter of a Poisson distribution. We defined *ŷ*_*i*_ for each training example as the prediction of the learning algorithm. This value corresponds to the mean of the predicted Poisson distribution.

The measure of the performance of the model is made through an objective function *O*(*θ*)= *L*(*θ*)+ Ω(*θ*) that sums the training loss *L* and the regularization term (penalty for complexity) Ω. The training loss to be minimized is then defined as the negative log-likelihood over the full set:

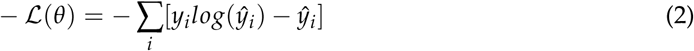

also known as the Poisson loss.

For the regularization term Ω, the complexity of the tree set was defined as

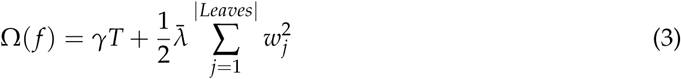

where *T* is the total number of leaves and *w*_*j*_ the score of leaf *j. γ* and 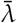 are two free parameters weighting the contribution of the two previous items in the objective function. For the sake of comparison with a related study Benjamin et al. [2017], we used the same values: *γ* = 0.4 and 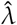 = 0.0. However, in the following section detailing the methods, we keep these two parameters as variables.

To minimize the objective function, the learning algorithm must find the optimal set of split values and the optimal set of leaf values for each tree. An efficient strategy is thus to optimize trees sequentially i.e. the input of a tree is the output of the previous tree. After optimizing the *t* - 1 trees, the prediction at tree *t* is 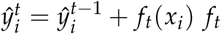 the function that maps the xi example onto the right leaf through the succession of tree partition.

By taking advantage of the fact that the same score is assigned to all the input data that fall into the same leaf, the objective function can be transformed from a sum over the training set to a sum over the leaves set:

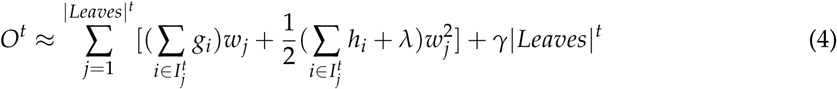

The index function *I*_*j*_ = {*i|f*(*x*_*i*_) = *w*_*j*_} maps each training point *x*_*i*_ to the corresponding leaf *j* while *g*_*i*_ and *h*_*i*_ are respectively the first order and second order derivatives of the loss function. In the case of Poisson regression, the *gi* and h_i_ are defined as:

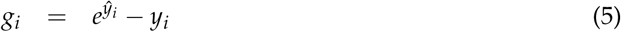

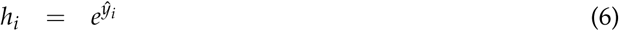

Finally, the sum of *w*_*j*_ and 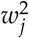 in equation 4 is quadratic, which allows us to compute the optimal 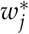 and the corresponding best objective value

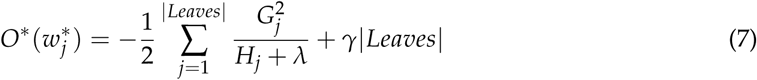

with *G*_*j*_ = Σ*i∊I_j_ g*_*i*_ and *H*_*j*_ = Σ*i∊I_j_* and *H*_*j*_ *h*_*i*_

The best tree structure is then found by sequentially splitting the features space, with each splitting position corresponding to the maximum gain:

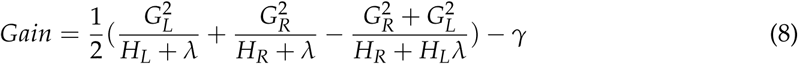

The gain for one split is a measure of fit improvement. It is the difference between the scores of the new leaves (subscripts *R,L*: right and left leaves, respectively) after the split and the score of the previous leaf. Details of the derivative steps and full explanations of the algorithm can be found in Chen and Guestrin [2016].

An example of the gradient boosted trees algorithm is shown in Fig. 1 for a non-linear tuning curve (blue curves Y). For each tree sequentially optimized (1,2 and 10 shown), the algorithm splits the tuning curve at different positions (X0, X1, X2, X3,…) and assigns a leaf score between each splits. By iterating this procedure, the predicted firing rate (black curves *Ŷ*) progressively converges to the actual firing rate.

**Figure 1:**
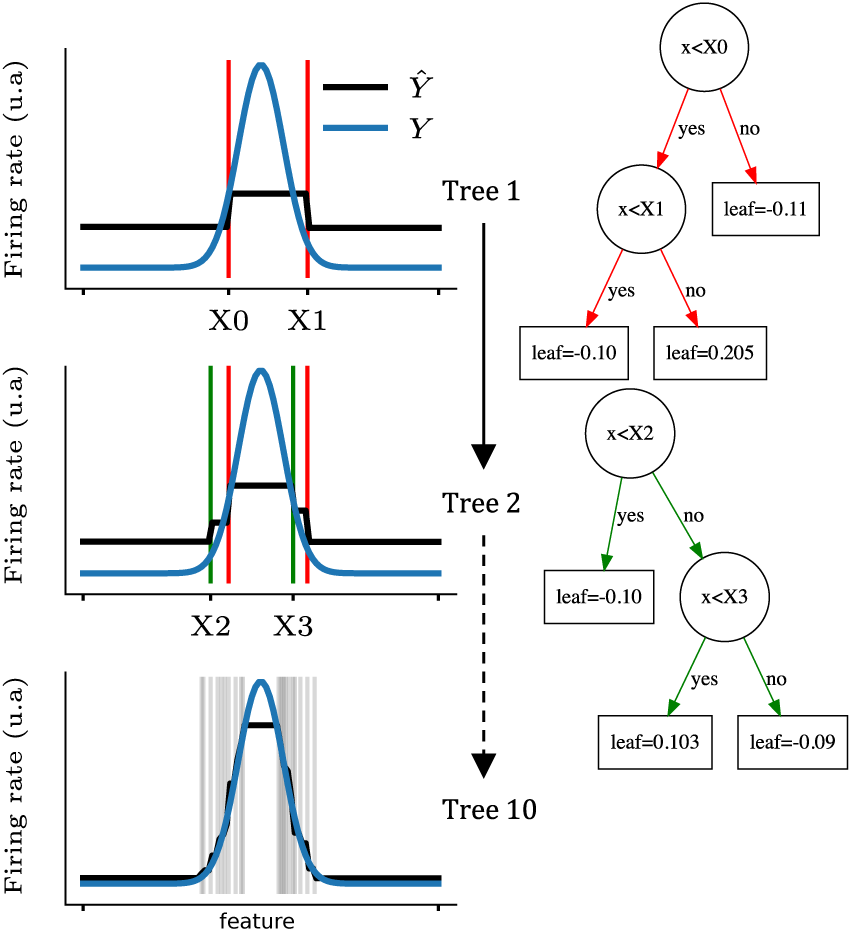
Predicting the firing rate of a cell with gradient boosted trees. Each row corresponds to the learning of one tree by the algorithm. The tuning curve is sequentially split as shown on the left figures (vertical lines; blue line displays the actual tuning curve and black lines correspond to the prediction). Thus, intervals between each pair of splits are assigned a different target value. The first two trees are shown on the right and the exact values of each leaf are indicated in the square boxes. Note that the predicted firing rates are the sum over all the leaves (i.e. the value of a single leaf can not be directly interpreted.

### Scoring function

To estimate the quality of a model, we used the pseudo-*R*^2^ score:

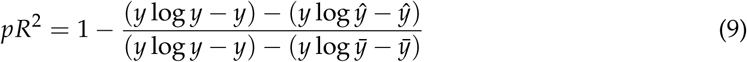

with *y* the target firing rate, *ŷ* the prediction, 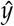 the mean firing rate [Cameron and Windmeijer, 1997]. A value of 1 indicates a perfect model that reproduces entirely the dataset while a value of 0 indicates a model that is no better than the average value of the training set.

To compute the pseudo-*R*^2^ score, the data set was divided into a training set and a test set, a procedure known as cross-validation, that prevents the model from over-fitting the training set. For all the predictions of firing rates, we used an 8-fold cross-validation, i.e the training set was divided into 8 discontinuous partitions with each one serving successively as the testing set. For each spiking activity predicted for one neuron, this procedure yields eight *pR*^2^ that were averaged. This mean *pR*^2^ served as a measure of performance of different techniques that were tested.

### Model comparison

In the present manuscript, we compare the prediction performance of XGB with three other methods. To this end, we computed the pseudo-*R*^2^ obtained with each method in an 8-fold cross-validation procedure. First, we tested a linear regression model between the animal’s HD and the binned spike trains. However, this method necessarily fails as the relation between the HD (an angular value) and the number of spikes emitted by HD cells is, in general, not linear. Therefore, we next linearized the HD by projecting the HD angular values on the first six harmonics of 2*pi* (called the 6^th^ order kernel in figure 2.B) and performed a linear regression with binned spike trains. Thus, a training point *x*_*i*_ corresponding to the direction *θ*_*i*_ is defined as a 12-dimensional input vector: *x*_*i*_ =[…, *cos(kθ*_*i*_), *sin(kθ*_*i*_),…] for k in [1,…, 6]. Finally, we tested a ‘model-based’ method: the tuning curve of a given HD neuron was computed from the training set and then used to predict the firing rate of the neuron in the test set.

**Figure 2:**
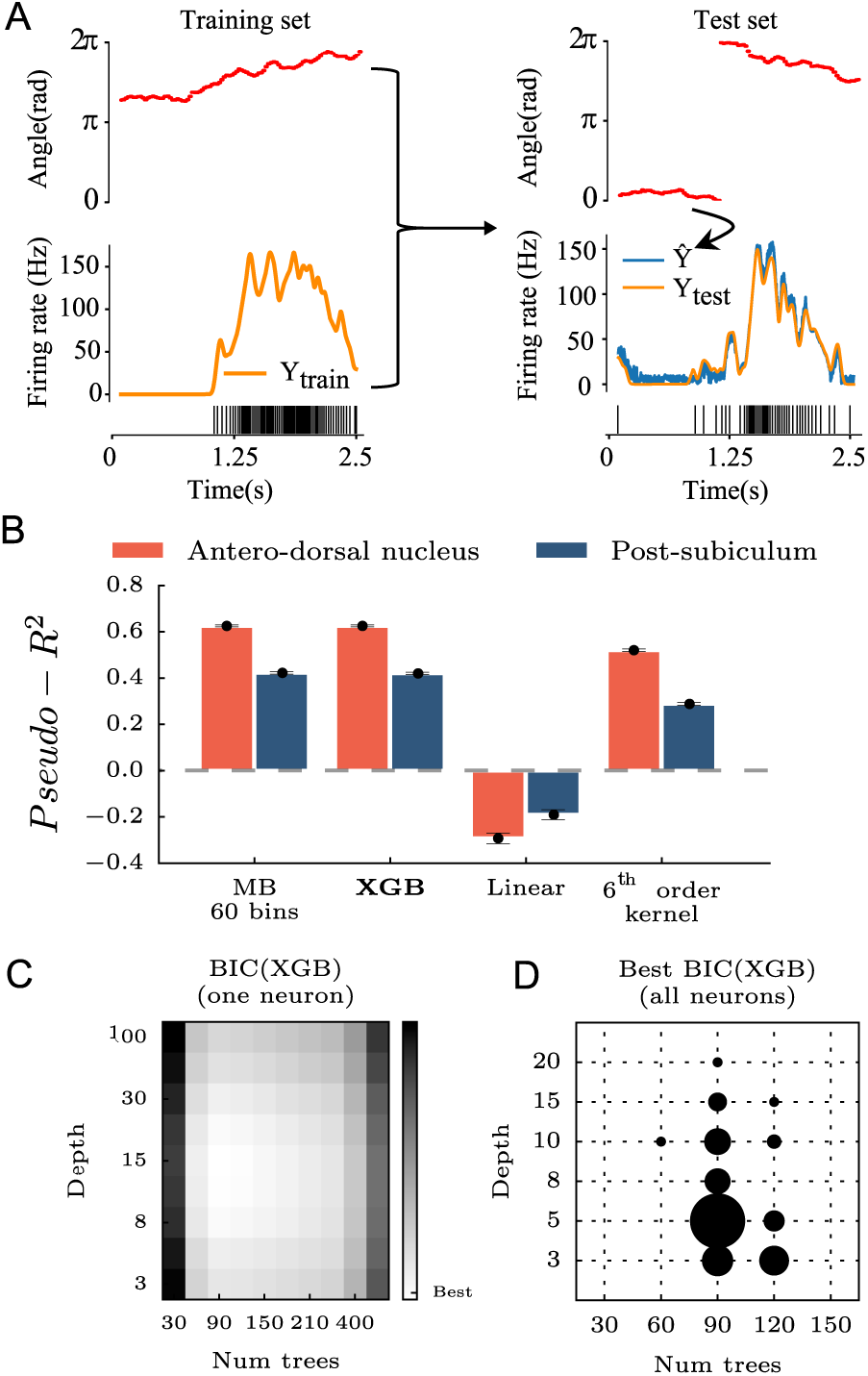
Comparing gradient boosted trees (XGB) with classical methods. **A** Using the angle as the input feature (red), the Machine Learning algorithm is trained to minimize the error in predicting the firing rate of one HD neuron over time (yellow, spiking activity below) during the training phase. For each angular position in the test set, the algorithm predicts a firing rate (blue curve). The score of the algorithm measures how close the prediction is to the real value. **B** Using an 8-fold cross-validation, XGB was compared to model-based tuning curves (MB) with 60 bins, a linear regression model and a linear regression model with preprocessing of the features i.e the first six harmonics of the angular direction of the head were used instead of the raw angle. Recordings from ADn and PoSub were used to benchmark each model. **C** To find the optimal number of trees and the optimal depth of XGB, a grid-search was performed for each neuron using the Bayesian Information Criterion (BIC). **D** Distribution of the set of optimal parameters for all neurons. Overall, a maximum number of 100 trees with a depth of 5 was used to learn and predict spiking activity as in **A**.

### Fisher Information

Fisher Information (FI) is directly related to the variance of the most optimal decoder and can be computed, under the assumption of a Poisson Process, directly from the tuning curve [Brunel and Nadal, 1998]. For recall, *FI*(*x*)=(*df /dx*)^2^/ *f* (*x*) with f(x) the firing rate at position *x* of the input feature. In practice, the Fisher Information was reduced to the squared slope of the line fitted between three successive bins of the tuning curve divided by the firing rate of the middle bin.

### Dataset

Neuronal recordings that are analyzed in this report were described in a previously published paper [Peyrache et al., 2015] and are available for download (https://crcns.org/data-sets/thalamus/th-1/). Briefly, multi-site silicon probes (Buzsaki32 and Buzsaki64 fom Neuronexus) were inserted over the antero-dorsal nucleus (ADn) of the thalamus in 7 mice. In three of these animals, a second probe was lowered to the post-subiculum (PoSub).

During the recording session, neurophysiological signals were acquired continuously at 20 kHz on a 256-channel Amplipex system (Szeged; 16-bit resolution, analog multiplexing). The wide-band signal was downsampled to 1.25 kHz and used as the local-field potential signal. To track the position of the animals in the open maze and in their home cage during rest epochs, two small light-emitting diodes (LEDs; 5-cm separation), mounted above the headstage, were recorded by a digital video camera at 30 frames per second. The LED locations were detected online and resampled at 39 Hz by the acquisition system. Spike sorting was performed semi-automatically, using KlustaKwik (http://klustakwik.sourceforge.net/). This was followed by manual adjustment of the waveform clusters using the software Klusters.

In animals implanted over the antero-dorsal nucleus, the thalamic probe was lowered until the first thalamic units could be detected on at least 2-3 shanks. The thalamic probe was then lowered by 70-140 *μ*m at the end of each session. In the animals implanted in both the thalamus and in the post-subiculum, the subicular probe was moved everyday once large HD cell ensembles were recorded from the thalamus. Thereafter, the thalamic probes were left at the same position for as long as the quality of the recordings remained high. They were subsequently adjusted to optimize the yield of HD cells. To prevent statistical bias of neuron sampling, we discarded sessions from analysis that were separated by less than 3 days during which the thalamic probe was not moved.

### Data analysis

In all analyses, spike trains were binned in 25 ms bins and smoothed with a 125 ms kernel, unless stated otherwise. The only exception is for decoding which was performed with bins of 200 ms. The animal’s HD was calculated by the relative orientation of two LEDs (blue and red) located on top of the head (see [Peyrache et al., 2015] for more details). The HD tuning curve of a neuron is the ratio between the histogram of spike counts as a function of HD (60 bins between 0 and 2*π*) and total time spent in each bin of HD. For a given angular bin *ϕ*_*i*_, the average firing rate is thus:

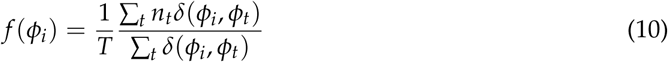

where *δ*(*ϕ*_*i*_, *ϕ*_*t*_) = 1 if, at time *t*, the angular HD *ϕ*_*t*_ is equal to *ϕ*_*i*_ (*δ*(*ϕ*_*i*_, *ϕ*_*t*_ otherwise), *n*_*t*_ the number of spikes counted in the *t*th time bin and T = 25ms (the time bin duration).

### Bayesian Decoding

The goal of Bayesian decoding in this study is to predict the HD of the animal given the spiking activity of recorded neurons. Let **n** = (*n*_1_, *n*_2_,…, *n*_*N*_) be the numbers of spikes fired by the HD neurons within a given time window (200 ms) and Φ the set of possible angular direction between 0 and 27*π*. The algorithm computes the probability *P*(Φ**n**) using the classical formula of conditional probability:

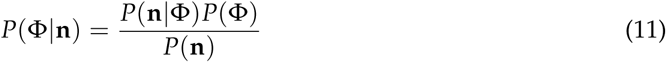

Assuming the statistical independence of HD neurons and the Poisson distributions of their spikes, the probability *P*(**n**Φ) can be evaluated as:

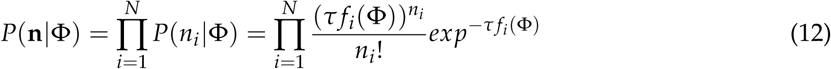

with *τ* the length of the time window and *f*_*i*_ (Φ) the average firing rate of cell *i* at position Φ. The full detail of the algorithm can be found in Zhang et al. [1998].

When using XGB for decoding the HD, we set the algorithm to do multiclass classification: the algorithm returns the predicted probabilities that population vector n (a vector of spike count of each neuron) belongs to each ‘class’ Φ =(*ϕ*_1_, *ϕ*_2_,…, *ϕ*_*k*_), i.e. 60 bins of HD. Briefly, learning of the decoder is achieved by minimizing the so-called ‘logarithmic loss’ computed as 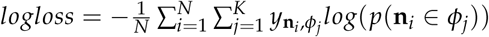 where *N* is the number of data points, *K* the number of classes (60 bins of HD in our case), *y*_**n**_*i*__, *ϕ*_*j*_ = 1 if the data point **n**_*i*_ is in class *ϕ*_*j*_ and 0 otherwise, and *p*(**n**_*i*_ ∈ *ϕ*_*j*_) is the predicted probability that observation **n**_*i*_ is in class *ϕ*_*j*_. Thus, a perfect classifier would have a null log loss (for each data point, there is one and only one class that has a probability p = 1 and that is correctly labeled, i.e. y = 1).

### Spiking network simulation

To attest the robustness of our analyses, the methods presented in this study were tested on an emulation of spiking neuronal ensembles using the Brian simulator Goodman and Brette [2009]. The network is composed of two layers of Poisson spiking neurons (*P*_*ADn*_ and *P*_*PoSub*_) and one layer of integrate-and-fire neurons (*I*_*PoSub*_). Poisson spiking neurons were individually parameterized by angular tuning curves. We used the actual HD of an exploration session (20 min) to generate a time-array of firing rate per neuron, at every time step of the simulation. Integrate-and-fire neurons follow a stochastic differential equation:

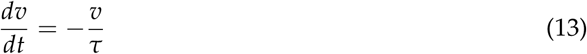

with the membrane time constant *τ* = 50ms for all simulations. We set the spiking voltage threshold *v* = 1 and after-spike reset to *v* = 0 and no refractory period.

The simulated integrate-and-fire neurons *I*_*PoSub*_, emulating spiking activity of observed PoSub neurons, received two sets of inputs. First, an input mimicking their actual tuning curve, each *I*_*PoSub*_ neuron receiving a connection from one *P*_*PoSub*_ neuron with a weight of 0.9. In other words, each integrate-and-fire *I*_*PoSub*_ neuron had a unique mirror Poisson spiking neuron in the *P*_*PoSub*_ layer that provides major driving input depending on the angular HD. The second set of synapses to *I*_*PoSub*_ were from a population mimicking ADn neurons, *P*_*ADn*_, with full connectivity (i.e. *I*_*PoSub*_ receives inputs from all *P*_*ADn*_ neurons). The weights of the connections from *P*_*ADn*_ units and a given *I*_*PoSub*_ neuron were parameterized by the angular distance between the preferred direction of the *I*_*ADn*_ and its pre-synaptic *P*_*ADn*_ neurons. More specifically, for two neurons *i* and *j* with respective preferred angular directions *ϕ*_*i*_ and *ϕ*_*j*_, the synaptic weight is defined as:

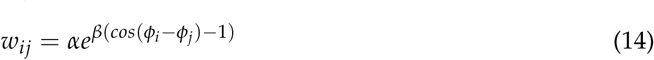

with *α* = 0.1 and *β* = 10.

### Code availability

The analyses presented in this report were run on Matlab (Mathworks, 2017) and Python. Code is available online in a raw form and as a Jupyter notebook to present some of the analyses (www.github.com/PeyracheLab/NeuroBoostedTrees). Gradient boosting was implemented with the XGBoost toolbox [Chen and Guestrin, 2016].

## Results

### Gradient boosted trees predict firing rates with raw features

We applied gradient boosted trees (XGB) to the prediction of spike counts from HD neurons recorded in ADn and PoSub (see Methods and Fig. 2.A for a full display of the training process). Since the HD signal is a well-characterized signal relative to the angular direction of the animal’s head, we compared the prediction of XGB with the output of the model-based (MB) tuning curve (that is, the firing rate expected from the HD of the animal knowing the tuning curve; see Fig. 2.B). The comparison shows that XGB reaches the same level of performance as MB for both ADn and PoSub. We then tested a generalized linear regression model with raw HD values or a 6^*th*^ order kernel. In the first case, the model learns only from the angular features *θ* ranging from 0 to 2*π*. In the second case, the model learns with all the k harmonics (*cosθ, sinθ,…, coskθ, sinkθ*). A 6^*th*^ order projection was used as it can fit the typical width of a HD cell tuning curve (approximatively 60 degrees at half peak). Not surprisingly, the simple linear model showed negative or null performances for both anatomical structures, because the relationship between a raw angular value and a binned spike train is unlikely linear (Fig. 2.B). Preprocessing of the angular feature (with the 6^*th*^ order kernel) increased the performance to the same levels as XGB and MB.

In comparison with XGB, linear models and MB are straightforward models in terms of numbers of free parameters. We thus performed a grid-search to find the optimal number of trees and depth of each tree to find the best estimate of the performance, measured by the *pseudo - R*^2^ (see Methods). A Bayesian Information Criterion (BIC) score (Fig. 2.C) was used to compare grid points. The BIC score was defined as *BIC(|Trees|, Depth*) = (|*Trees*|+ *Depth*)*log*(*n*)- *2log(L)* with *n* the number of time steps in the data training set and L the likelihood of the model. By penalizing more complex models using this approach, we found that 100 trees with a maximal depth of 5 were sufficient to predict spike trains for all neurons (Fig. 2.C,D).

### Decoding of brain signals

Once the relationship between a behavioral feature and spiking activity has been learned, XGB can be used to decode the internal representation of this feature based on population spiking activity. We thus tested its performance on the decoding of the HD signal distributed over population of HD cells. To this end, spiking activity was binned in 200ms windows and XGB was trained and compared to a Bayesian decoding method, a technique widely used for such tasks [Peyrache et al., 2015, Zhang et al., 1998], that predicts the probability of having a particular HD at each time step based on the instantaneous spike count in the population. For both algorithms, 60 angular bins were used to predict the HD. We parametrized the gradient boosted trees to use the multi-class log-loss that outputs a probability of being in a certain class or not (see Methods).

We decoded the HD signal in sessions that contained more than 7 neurons in both ADn and PoSub (n=5 sessions, two animals). An example of 30 second decoding for XGB is shown in Fig. S1.A. Gradient boosted trees and Bayesian decoding show similar performances when using ADn activity as a feature while gradient boosted trees slightly outperforms Bayesian decoding for PoSub activity (Fig. S1.B). In addition, we observed that the decoding of the HD from ADn firing rate outperforms the decoding of the head direction using PoSub activity. This observation was consistent for both methods.

### Information content of the feature space is revealed by data splitting

Gradient boosting, as most Machine Learning tools, can be considered a black box that achieves high levels of performance while the particular details of the learning procedure remain unknown. However, it is possible to retrieve the thresholds at which trees split the data to predict the target output (as shown in Fig. 1). In the case of HD cells, whose firing was directly predicted from the HD of the animals, splits concentrated on HD values where the tuning curves were the steepest (see examples of figure 3.A). In fact, the density of splits is strongly correlated with the Fisher Information (Fig. 3.B), a measure that is related, but not equal, to tuning curve steepness and that estimates the variance of an optimal decoder [Brunel and Nadal, 1998].

**Figure 3:**
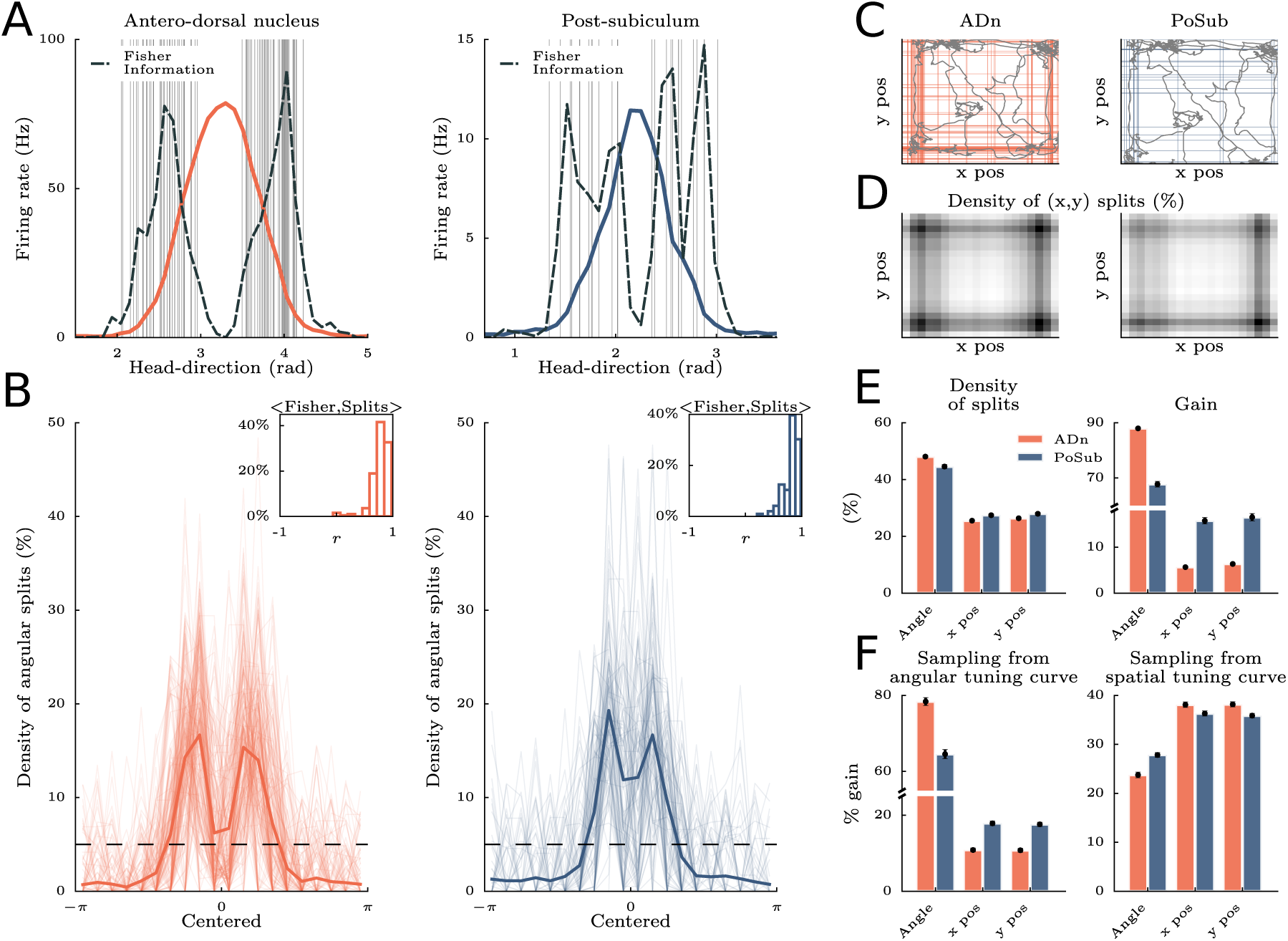
Segmentation of behavioral features to predict neuronal spiking. **A** Tuning-curve splitting for one neuron of the antero-dorsal nucleus (ADn) and one neuron of the post-subiculum (PoSub). Each vertical gray line is a split from the gradient boosted trees used to predict firing rate. Dashed black lines indicate Fisher Information (computed from the tuning curves). **B** Density of angular splits for ADn and PoSub for all the neurons, and average (thick line). Splits positions were realigned relative to the peak of the tuning curve. Horizontal dashed lines display chance levels. Insets show the distribution of correlation coefficients between Fisher Information and density of splits. **C** Using x and y coordinates of the animal in the environment as additional input features of the algorithm. Colored lines indicate spatial positions of splits along x and y. Gray lines indicate a short segment of the trajectory of the animal during the example session. **D** Density of splits for x and y position features for all neurons. The highest density is shown in black. **E** Left. proportion of splits for the three input features (head direction, x position and y position for ADn and PoSub. Right. Mean gain value for the three input features (head direction, x position and y position for ADn and PoSub) **F** Same as **E** for the gain value except that the firing rate for each neuron was generated from the angular tuning curve (left) or the spatial tuning curve (right)

Many neurons of the brain’s navigation system exhibit correlates to more than one behavioral parameters, for example HD and place [Cacucci et al., 2004, Peyrache et al., 2017, Sargolini et al., 2006]. We thus predicted spike trains based on the three observed behavioral features, assuming they were independent: x and y positions of the animal randomly foraging in the environment, as well as the HD. We thus increased the feature space and dissected the resulting splitting distribution of the gradient boosted trees. In average, the density of splits along the (*x, y*) coordinates was the highest in the corner of the environment (Fig. 3.C and D) where animals naturally spend a large amount of time. Analysis of the distribution of splits reveals that the HD feature was more segmented than the (*x, y*) coordinates for both ADn and PoSub (Fig. 3.E, *left*), showing that HD neurons in both ADn and PoSub are primarily driven by HD. Nevertheless, we observed that the proportion of positive splits relative to angular splits was slightly higher for PoSub when compared to ADn.

One potential issue with this approach is that training a large number of trees overfits the learning procedure: it is optimal for decoding performance but not necessarily for the interpretability of the tree structure. To best explain the contribution of various features to the spiking activity, it is sometimes more suited to concentrate on the structure of a smaller number of trees, and examine the ‘gain’ of each feature when training the first trees. In fact, the average gain (see equation 8) for each feature decreases exponentially as the number of trees increases (Fig. S2). In addition, we found that random features were also more split as the number of trees increased (see Fig. S3). For all these reasons, we restricted our analyses to the characteristic decay constant of the gain as a function of number of trees (see Fig. S2), i.e. 30 trees with a depth of 2.

Shifting from split density to gain analysis, we thus demonstrate that the gain of spatial features (x and y position) was approximatively three times higher for PoSub neurons compared to ADn neurons (Fig. 3.E *right*), in agreement with previous studies that employed model-based methods (i.e. that assumed various properties of spike trains and sampling of the feature space) [Cacucci et al., 2004, Peyrache et al., 2017]. To assess that the advantage of angular information over spatial information was not caused by a difference in the trajectories of the animals (i.e sub-sampling of some portions of the 2 dimensional space), we generated, for each neuron, artificial spike trains sampled from either the angular or spatial tuning curves. In the case of angular tuning curve sampling, we found qualitatively the same gains for PoSub and ADn neurons (Fig. 3.F, *left*). When sampling the spatial tuning curves to generate artificial spike trains, gains for spatial features were higher than for HD, as expected (Fig. 3.F, *right*). However, the difference with HD gains was small, and the gains were not different for ADn and PoSub neurons, indicating that the place fields of these two classes of neurons do not convey much spatial information. Thus, we concluded that XGB, when used appropriately, is an efficient method for determining the relative contribution of various features to a series of spike trains.

### Performances of peer-prediction

Brain functions arise from the communication of neurons with their peers in local and downstream networks. However, how these interactions take place remains largely unknown. With this question in mind, we thus applied XGB to neuronal peer-prediction, that is learning to estimate the spiking activity of one neuron as a function of the activity of a population of other, presumably anatomically-related neurons (Harris et al. [2003], Peyrache et al. [2015], Pillow et al. [2008]). For each session that contained at least 7 neurons in both ADn and PoSub, the model learned all possible group combinations (ADn->ADn, PoSub->ADn, PoSub->PoSub, ADn->PoSub). This learning was performed with no spike history, i.e. the bins used as features were synchronous to the bin predicted. For intra-group prediction, the target neuron was removed from the pool of feature neurons. Tested during wake, REM and non-REM sleep, we found that peer-prediction had the highest prediction score between ADn neurons and the lowest score between PoSub neurons (Fig. 4.A). Inter-group predictions were similar. In all cases, scores during non-REM sleep were systematically lower than during wakefulness and REM, in agreement with previous analysis of peer-prediction in thalamo-cortical assemblies Peyrache et al. [2015].

**Figure 4:**
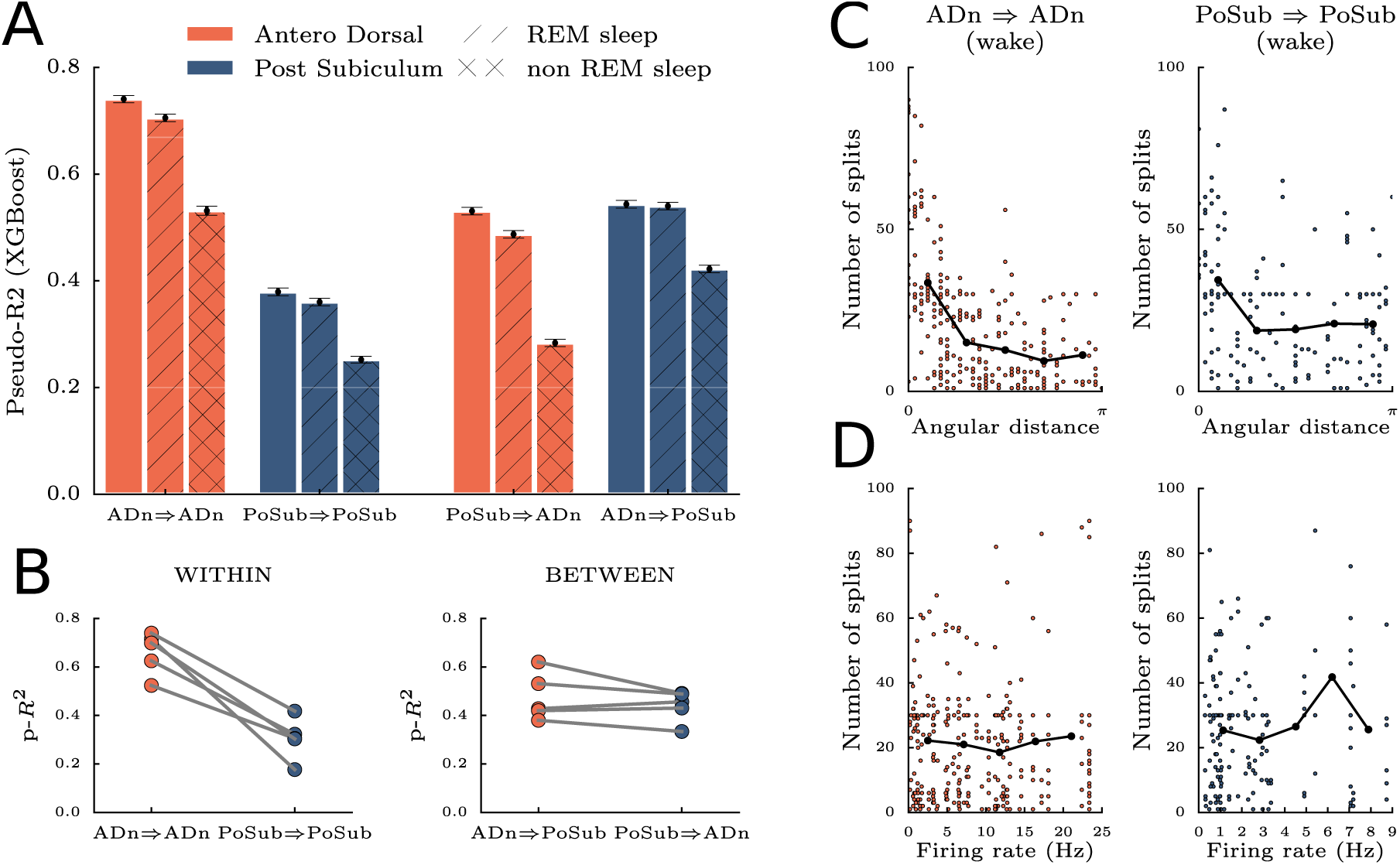
Peer-prediction between ADn and PoSub. **A** Two conditions were tested: prediction between neurons of the same population (ADn⇒ADn and PoSub⇒PoSub) and prediction using neurons of the other population (PoSub⇒ADn and ADn⇒PoSub). Only sessions with at least 7 neurons in each population were included (2 animals). Peer-prediction was then tested during wake (plain bars), REM sleep (dashed bars) and non-REM sleep (crossed bars) episodes. **B** To rule out the possibility that the difference in scores resulted from uneven number of recorded neurons, the score were recomputed using an equal number of neurons in each population (i.e by randomly selecting neurons within the largest group). **C** Number of splits of one feature neuron given its angular distance with the target neuron. **D** Number of splits given the mean firing rate of the feature neuron. Despite firing rate differences, all features neurons contributed.

An uneven number of feature neurons is a potential confound in peer-prediction analysis. The prediction process was thus repeated by equalizing the number of neurons in both structures and it yielded scores similar to the original analysis (Fig. 4.B). The activity within ADn is therefore more predictable than in the PoSub.

To best capture the statistical dependencies between spikes trains, we focused on a gain analysis (i.e. from the branching structure resulting from learning on only 30 trees with a depth of 2) and we found that the angular distance was a weak predictor of the split density for both ADn and PoSub (Fig. 4.C). In others words, gradient boosted trees tend to split preferentially, yet mildly, the instantaneous firing rate of feature neurons that have a preferred direction closer to the target neuron. More surprisingly, we found no correlation between the mean firing rate of neurons and the density of splits (Fig. 4.D). Feature data from neurons with high firing rates are characterized by a wider range of values to be split, yet, this does not lead to increased splitting. Thus, all neurons contributed to the prediction of the activity of another neuron despite each idiosyncratic spiking activity.

### Peer-prediction reveals the directionality of information flow across brain structures

While the HD signal is aligned with the actual heading of the awake animal in the PoSub, the spiking of HD cells in the ADn are best explained by the future heading of the animal, by about 10-50 ms Blair and Sharp [1995], Taube and Muller [1998]. This finding suggests that the neuronal activity in the ADn should lead PoSub spiking at least during wakefulness, perhaps in all brain states. We thus tested the ability of XGB to reveal the temporal constraints of neuronal communication across brain areas compared to the classical cross-correlation of spike train pairs. One issue with linear cross-correlation analysis is that it is dominated by the slow dynamics of the underlying signal and, while the HD signal has comparable dynamics during wake and REM sleep, it is accelerated during non-REM sleep [Peyrache et al., 2015]. During wakefulness and REM, cross-correlations do not reveal clear bias in the temporal organization of the ADn to PoSub communication. Furthermore, a Principal Component Analysis of the cross-correlograms reveals that, overall, cross-correlograms of thalamo-cortical pairs of neurons are rather good indicators of the ongoing brain state (Fig. 5.A, *left*). Finally, the sign of the correlation between HD neurons depend, in all brain states, on the angular difference of their preferred direction [Peyrache et al., 2015]. Thus, cross-correlograms are in average flat (Fig. 5.A, *right*), and the overall effect can only be captured by the study of the variance of the cross-correlograms. The variance of the corr-correlograms shows a slight biases for negative latencies from ADn to PoSub (insets in Fig. 5.A, right), but, again, the variance profile (and thus its resolution) depends on the brain states.

**Figure 5:**
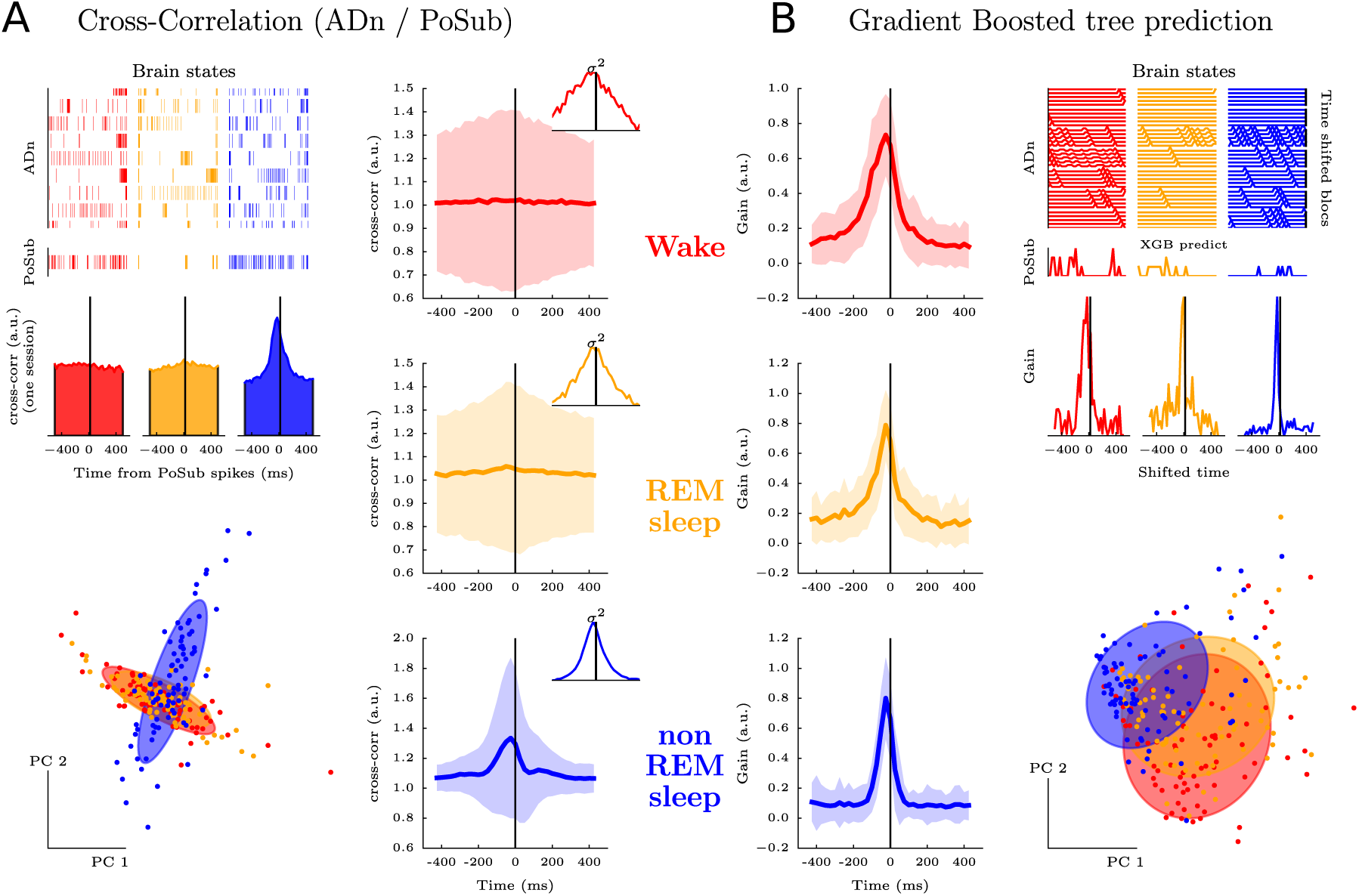
Temporal coordination between ADn and PoSub is preserved across brain states, as revealed by gradient boosted trees but not cross-correlation. **A** Cross-correlation between spike trains of ADn and PoSub neurons during the Wake, REM sleep and non-REM sleep (respectively in red, yellow and blue). Top left, examples of spiking activity for the three brain states; middle left, one-session example of an average cross-correlation between an ensemble of ADn neurons and one PoSub neuron; bottom left first two dimensions of a PCA performed on all cross-correlations, across all three brain states and neurons of PoSub. Colored circles are the best Gaussian fit for each state, showing that Wake and REM sleep yield qualitatively similar cross-correlations, but not non-REM sleep. Right, averaged (± s.d.) cross-correlation for each state. The insets shows the variance. **B** Gradient boosted tree prediction of PoSub firing from the activity of ADn neuron ensembles at successive past to future time steps during Wake, REM sleep and non-REM sleep. Top right, example instantaneous firing rates an ADn neuronal ensemble (shifted at various positive and negative lags) and a PoSub neuron; middle right, prediction gain for the example session; left, the gain of the algorithm was maximal around 25 ms before PoSub spikes (vertical black lines) for wake and both sleep stages; bottom right PCA of all resulting gain, across brain states. The best Gaussian fits of each state (as in panel A) are now overlapping.

Can XGB reveal the temporal component of neuronal communication across brain areas? To investigate this question, XBG was run for peer-prediction of individual PoSub neurons from multiple copies of ADn population activity at various time-lags. In other words, the model learned the relationship between the firing rates of feature neurons from time *t* - *T* to *t* + *T* (in Fig. 4.A, the model had access only to time *t*). A graphical explanation of this procedure is shown in Fig. S4. Using only raw, unsmoothed spike counts, we found that the gain (the number of splits multiplied by the mean gain) was maximal at −25 ms when predicting PoSub firing rate with ADn activity (Fig. 5.B), in agreement with the anticipation delay of ADn HD neurons [Blair and Sharp, 1995, Taube and Muller, 1998]. The distribution of transmission delays was only weakly dependent on brain states, suggesting a hard-wired, internally organized circuit (Fig. 5.B) [Peyrache et al., 2015].

### Robustness of gradient boosted trees to detect delay of transmission is asserted by a spiking network

**Figure 6:**
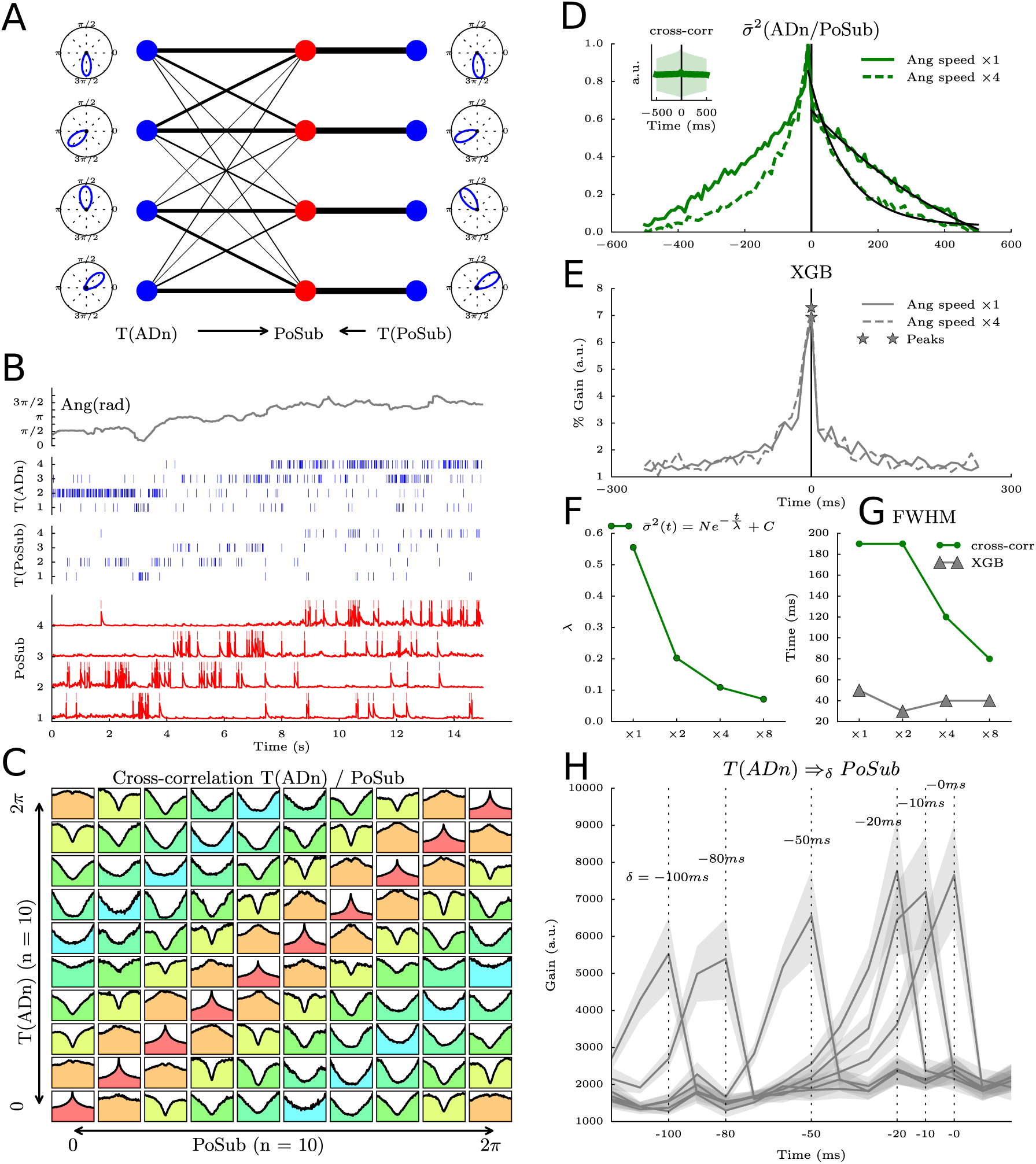
Spiking network simulations reveal the robustness of gradient boosted trees to detect transmission delays independent of feature dynamics. **A** The layer of PoSub integrate-and-fire neurons (red dots) receives one-to-one input from a mirrored layer of neurons which determines their primary angular tuning curves (right T(PoSub) in blue dots) and inputs from a layer of ADn neurons (left T(ADn) in blue dots) with full connectivity. The synaptic weight from T(ADn) to PoSub is proportional to the angular difference between the respective tuning curves of ADn neurons and PoSub neurons. **B** Simulation of 15 s of data. Top row, real HD value of one animal. Middle, raster of spiking activity of T(ADn) (top) and T(PoSub) (bottom). Bottom, membrane potential of the PoSub neurons. **C** Cross-correlograms between the spiking activity of 10 T(ADn) neurons and 10 PoSub neurons sorted according to the angular peak of their tuning curves. The angular difference between the preferred firing directions is color-coded (0 in red, *π* in blue). **D** Centered standard deviation of the cross-correlograms at normal (full green line) and accelerated angular speed (dashed green line). Synaptic transmission is set at 0 in these simulations. Black lines show the best exponential fits. **E** Same as **D**, but using XGB peer prediction of PoSub spiking activity from T(ADn) activity. Note that the distribution peaks at 0 ms for both angular speeds. **F** Characteristic time decays of the cross-correlogram exponential fits as a function of angular speed. **G** Full width at half maximum (FWHM) of cross-correlograms and XGB learning gain as a function of angular velocity. **G** XGB gains as a function of synaptic delays of transmission between T(ADn) and PoSub.

To assess that gradient boosting can determine temporal shifts between spike trains of neurons in vivo, independent of brain-state dynamics (i.e. feature dynamics), we further tested the methods with smulations of spiking networks Goodman and Brette [2009]. We first sought to replicate the temporal delay between ADn and PoSub shown in Fig. 6.A (see Methods). To this end, we used HD tuning curves and the animal’s HD to generate series of spike trains in an artificial population of ADn and PoSub neurons. Those neurons are Poisson spiking neurons parameterized at each time step only by the instantaneous firing rate read from the angular tuning curves, thus referred to as T(ADn) and T(PoSub). We then modeled a population of PoSub integrate-and-fire neurons that receive one-to-one inputs with a fixed weight from T(PoSub) and multiple inputs from T(ADn) with synaptic weights inversely proportional to the angular distance (Fig. 6.A). As shown in Fig.6.B for four neurons of each layer, the neurons of PoSub fired whenever the animal’s HD crossed their angular tuning curves. To demonstrate that PoSub neurons integrate information that is related to the tuning curves of T(ADn), we showed the cross-correlation between each pair of neurons from the two layers, sorted by their preferred angular direction (Fig.6.C). As with cross-correlations of pairs of real HD neurons Peyrache et al. [2015], pairs of HD neurons with overlapping tuning curves show positive correlations (i.e. peaks in the cross-correlgrams) and pairs of opposite preferred directions show negative correlation (i.e. dip in the cross-correlograms). As expected, the average cross-correlogram is flat (inset in Fig.6.D).

To reproduce the observation that the temporal width of cross-correlations was smaller for non-REM sleep than for REM sleep and wake, we gradually changed the speed of the animal’s HD in input. As expected, the temporal width of cross-correlations was primarily driven by the feature dynamics as shown in Fig.6.D for an angular speed accelerated four times. When doing peer-prediction with XGB as in Fig.5.B, we observed that the prediction of time lag remained qualitatively the same when the angular speed was accelerated four times (Fig.6.E). We quantified the decrease of temporal width in the cross-correlogram for four different speeds with an exponential decay fit (Fig.6.F) and the full width at half maximum (FWHM, Fig.6.G). In contrast, the FWHM resulting from XGB remained constant across conditions (Fig.6.G).

Could the model confirm that XBG accurately tracks synaptic delays? Even when synaptic transmission delay was set at 0 ms, the variance of the cross-correlograms was slightly shifted at negative time lags (Fig.6.D), unlike the XGB gain (Fig.6.E). This suggests that linear cross-correlations, but not XGB, is biased by the integration time constant of the post-synaptic neuron. In addition, we observed that changing the intrinsic transmission delay was fully captured by XGB (Fig.6.H). In conclusion, applying gradient boosted trees on neuronal ensembles reveals intrinsic temporal organization of the circuit, independent of the brain-state specific dynamics of the underlying features.

## Discussion

We show how non-linear encoders are versatile and useful tools to study neuronal data in relation to behavior and brain states. More specifically, we found using these methods that, in the HD system, the thalamus temporally leads the cortex during wakefulness and sleep, suggesting a bottom-up transmission of signal irrespective of the brain state.

While classical tools aim to provide interpretation of the data by investigating the predictability of a particular model of neuronal function, we show that gradient boosted trees [Chen and Guestrin, 2016, Friedman, 2001], a supervised learning technique commonly used in various fields of data mining, equals, if not outperforms other classes of Machine Learning models [Burges, 2010, Li, 2012]. This performance was achieved by a direct fit of raw behavioral or neuronal data to the targeted spike trains, with no explicit prior on a cell’s response (e.g. a tuning curve, or a model of mixed-selectivity to a set of variables). We report optimal parameters and detailed methods to study neuronal response and dynamics as a function of behavior or endogenous processes (e.g. the neuronal peer network). Furthermore, we show that the resulting tree structure, after learning of the data, can be itself analyzed to reveal important properties of the neuronal networks.

### Learning neuronal firing in relation to behavioral data: performance and optimal parameters

We first sought to validate the approach of learning a predictive model of spike trains from behavioral data with a decision tree learning algorithm that does not include a predefined model of the training set. To this end, we analyzed a dataset of HD cells [Peyrache et al., 2015, Taube et al., 1990], whose firing in relation to behavior is among the best characterized signals in the mammalian nervous system. We demonstrated first that gradient boosted trees predicts the firing of the neurons with high accuracy by establishing a direct correspondence between the raw behavioral data (in this case the HD angular value alone) and the instantaneous spiking of the neurons (Fig. 2). Using a Generalized Linear Model to regress the spiking activity of a neuron on raw behavioral data, such as the HD angular values, necessarily fails as this relationship is not generally linear. It is thus necessary to project the raw data on a set of orthogonal functions that linearize the inputs. Therefore, we used a basis of trigonometric functions up to the 6^*th*^ order that can, in theory, capture the typical width of a HD cell tuning curve (approximatively 60 degree width). In this case, the prediction performance was similar to XGB. The same type of transformation has been applied previously, for example Zernicke’s polynomials for position values in a circular environment Acharya et al. [2016]. However, it is clear from these two examples that one major strength of XGB is to generalize prediction to all possible behavioral data (e.g. not depending on the particular shape of an environment for position data). Finally, the performance of XGB was similar to a model-based approach (i.e. prediction of the firing rate on test data based on the tuning curve of the training set). This is not surprising for a class of neurons whose spiking activity is explained so well by an experimentally tractable signal. However, in general, tuning curves for even well defined neuronal responses explain actual spike trains only partially and XGB may well capture previously undetermined sources of variance.

Although XGB can be viewed as a model-free technique that does not assume any particular statistics or generative model of the input data, the procedure still depends on a limited set of free parameters that need to be tuned for optimal performance. To facilitate the use of this classifier for future studies and assure reproducibility of analyses across laboratories, we systematically explored the parameter space for depth and number of trees for spike train prediction. When computing prediction performance (measured by the pseudo-*R*^2^), we found that minima were well localized, for all neurons, using the BIC score that penalizes over-complex models. More specifically, we show how the use of multiple trees (approximatively 100), each limited in depth (typically five branching), was an optimal choice of parameters. Importantly, these optimal parameters did not seem to depend on a neuron’s intrinsic parameters (e.g. firing rates) and there was no obvious trade-off between tree depth and number of trees (the two optimal values were independently distributed across neurons).

### Interpreting the structure of the gradient boosted trees

While the structure of a multi-layered neural nets (or other forms of deep architecture) after learning the classification of a dataset is notoriously unanalyzable [Mikolov et al., 2013, Szegedy et al., 2013, Zeiler and Fergus, 2014], we show how the branching of the decision tree may be highly informative on how input data are matched to their output targets. The density of splits (or branching) across the series of trees was maximal in the range of inputs where firing rates vary the most. This could be interpreted as a maximum data splitting around the maxima of Fisher Information which is, for a Poisson process, directly related to the change in firing rate as a function of stimulus value, that is when spike trains are most informative about the encoded signal [Averbeck et al., 2006, Brunel and Nadal, 1998]. Although the relationship between tree branching and Fisher Information is, in our study, purely empirical, it is interesting to show, again, that unraveling the tree structure allows the understanding of how the data are learned by gradient boosting.

In the case of neuronal peer-prediction, analyzing the structure of gradient boosted trees presents the advantage that all kinds of neuronal interactions (positive, negative, linear or monotonically non-linear) yield comparable estimates (when quantifying split density or gain). This is in contrast with classical correlation analyses of individual spike trains relative to a population of peers that may be hard to interpret in certain cases where these interactions are both negative and positive ([Okun et al., 2015, Renart et al., 2010]. As we show here, linear correlations are also directly affected by the ongoing brain dynamics. In addition, fitting spiking data to maximum entropy (i.e. Ising) models have revealed that linear correlations may not indicate the true coor-dination between spike trains [Cocco et al., 2009, Schneidman et al., 2006]. The analysis of tree branching provides an estimate of the statistical dependencies between spike trains, independent of the underlying type of interaction and without assuming a particular transfer function for the target neuron [Harris et al., 2003]. The nature of neuronal coordination as observed from spike trains is still debated, for example in the hippocampus [Chadwick et al., 2015], and unbiased, model-free methods may be highly informative on the nature of the actual statistical dependencies between neurons.

We also report an optimal range of tree number that should be used for split analysis when regressing spike trains on behavioral features or the activity of other neurons (Figs. 3 – 5). ‘Learning gain’ decreases exponentially with the number of trees (Fig. 7). Using less trees (typically 30 with a depth off 2) allows the estimation of how different features contribute to the output target, at the expense of prediction and decoding performance (which are best estimated with approximatively 100 trees with a depth of 5, see above). In contrast, fitting the data on too many trees leads to overfitting and should be avoided. Overall, readers interested in using this technique should bear in mind that meaningful information about the dataset can sometimes be overshadowed by high split density. In such cases, it is of best interest to reduce the number of trees and to ensure that the average gain for splits is large enough.

**Figure 7:**
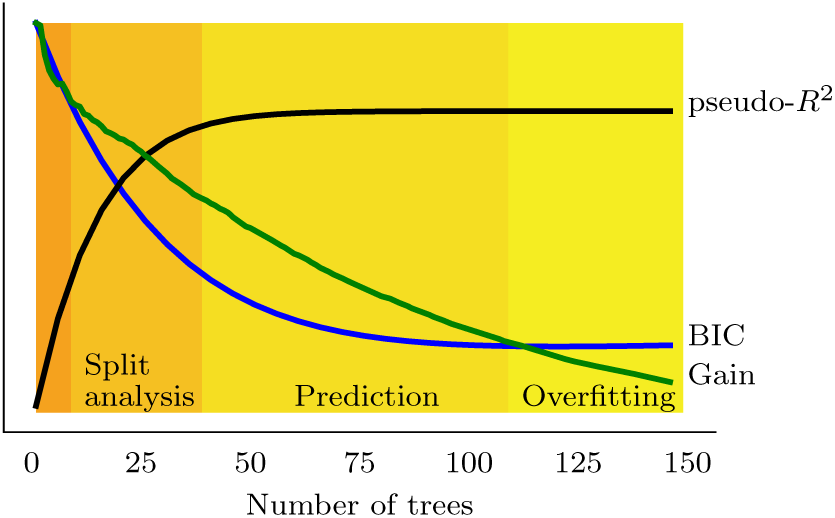
Split analysis and optimal data prediction lies within different ranges of tree numbers (for a fixed tree depth). Thus, the use of gradient boosted trees requires a careful tuning of the parameters of the algorithm depending on the question (interpretability of the structure versus prediction and decoding of the signal).

### Measuring the contribution of multiple behavioral variables

A large class of neurons in the brain are modulated by several dimensions of incoming stimuli [Finkelstein et al., 2015, Hardcastle et al., 2017, Rigotti et al., 2013, Sargolini et al., 2006], a property referred to as mixed-selectivity. Untangling the different contributions is sometimes challenging and gradient boosted trees offer a rapid and unequivocal approach to address this issue Benjamin et al. [2017], Truccolo and Donoghue [2007]. In fact, there is no intrinsic limit to the dimensionality of the inputs that can be learned. To further test this technique, we regressed spike trains of HD cells on spatial position, as well as on HD data. In line with previous reports [Cacucci et al., 2004, Peyrache et al., 2017], the HD cells of the PoSub correlated also with spatial factors while in the ADn, neurons coded mostly for the HD (Fig. 3). XGB thus enables to rapidly explore the correlates of spike trains to measurements of external or internal variables of the system.

### Prediction of feed-forward activation in a thalamo-cortical network in vivo

Investigation of neuronal dynamics does not always entail the regression of spiking data to variables of the experiments. Many studies have focused on the spatio-temporal coordination of neuronal networks in vivo, independent of any behaviorally-related processing ([Luczak et al., 2007, Okun et al., 2015, Peyrache et al., 2012]. In fact, the characterization of signal transmission between brain areas remains one of the most complex challenges of neuroscience as it first requires the recording of such data in vivo as well as the establishment of a proper model of interaction to determine the parameters of spike transmission (e.g. conduction delay and post-synaptic integration time).

Here we used data from the HD thalamo-cortical network [Peyrache et al., 2015] with simultaneous recording of PoSub and ADn. It allowed us to demonstrate a temporally-shifted relationship from ADn to PoSub. More precisely, we used gradient boosted trees to predict PoSub HD cell firing activity based on the ensemble spike trains of the HD cells of the ADn, at various time-lags between the two series of spike trains. PoSub spiking was mostly dependent on ADn activity in preceding time bins (in average 25ms), thus indicating a likely feed-forward pathway. First, this replicates the findings that the HD signal of ADn neurons precedes the actual HD by about 25 ms [Blair and Sharp, 1995, Goodridge and Taube, 1997, Taube and Muller, 1998]. Second, the temporal asymmetry in the prediction of cortical spiking relative to thalamic activity was preserved during sleep, both during REM and non-REM, and it therefore indicates that this differential temporal coding likely emerges from intrinsic wiring and dynamics. This confirms anatomical studies, as well as examination of putative synaptic interaction between neurons in this pathway Peyrache et al. [2015].

The robustness of this approach was validated by the analysis of artificially generated spike trains, drawn from actual tuning curves and in which different input feature dynamics (in our case, angular head velocity, or ‘virtual’ angular speed during sleep), transmission delays, and integration time constant were explored. This study confirmed the results of in vivo data: unlike linear cross-correlations, gradient boosting reveals temporal organization of spiking irrespective of the dynamics of the inputs and accurately extract, in all conditions, a delay introduced between spike trains (Fig. 6). Furthermore, while PoSub integration time constant alone results in temporarily shifted cross-correlograms between ADn and PoSub simulated spike trains, gradient boosting captures only the synaptic transmission delay.

### Gradient boosted trees match Bayesian decoding in performance

Neurons convey information about external parameters, and it should thus be possible to decode these signals from population activity. The best examples are the demonstrations that position can be estimated from ensembles of hippocampal place cells during exploration and ‘imagination’ of future paths [Johnson and Redish, 2007, Pfeiffer and Foster, 2013, Wilson and McNaughton, 1993] as well as the HD signal during wakefulness Johnson et al. [2005] and sleep Peyrache et al. [2015]. Decoding of neuronal signals has also been widely studied in the context of brain machine interface [Laubach et al., 2000].

Bayesian decoding is the tool of reference to estimate a signal from ensembles of neurons. In fact, it computes the probability distribution of a particular signal given the tuning curves of the neurons and the instantaneous spike counts in the neuronal population. This technique generally assumes that spike counts are drawn from Poisson processes and that neurons are independent from each other (Zhang et al. [1998]). Here we have compared the performance of Bayesian decoders and gradient boosted trees for decoding angular values based on the activity of either ADn or PoSub neuronal ensembles (Fig.S1). We found that gradient boosted trees matched Bayesian decoding when using ADn neurons but were slightly better with PoSub activity. As emphasized in this report (Fig. 4.E and F), PoSub activity does not encode only the HD but also spatial information about the location of the animal. In case of mixed-selectivity signals, a model-free technique such as gradient boosted trees is less impaired at predicting an external variable compared to the classical method of Bayesian decoding.

### Potential for neuroscience and future work

The potential of these methods to unravel the dynamics of biological neuronal networks is tremendous and will be the scope of further studies. For instance, tracking synaptic transmission in pairwise spike trains [Barthó et al., 2004], uncoupling the phase-locking of neuronal spiking to concomitant and nested brain oscillations [Belluscio et al., 2012, Tort et al., 2009], and determining the nature of the coordination in neuronal populations in relation to behavior [Chadwick et al., 2015, Harris et al., 2003] are examples of the current challenges of data analysis in systems neuroscience. In addition, future improvements of brain-machine interface will require the development of reliable and robust tools to decode neuronal activity [Andersen et al., 2004, Lebedev and Nicolelis, 2006].

In summary, gradient boosted trees methods are potentially helpful tools to explore a dataset and make a prediction on the underlying biological processes which, in turn, can be tested with more classical methods. They may also be used to decode signals for closed-loops experiments and brain-machine interface in animals or humans. Finally, these methods open avenues for the study of neuronal data, in general, as the branching of the tree structure can be analyzed as a ‘proxy’ of the biological system itself.

---

**Figure S1:**
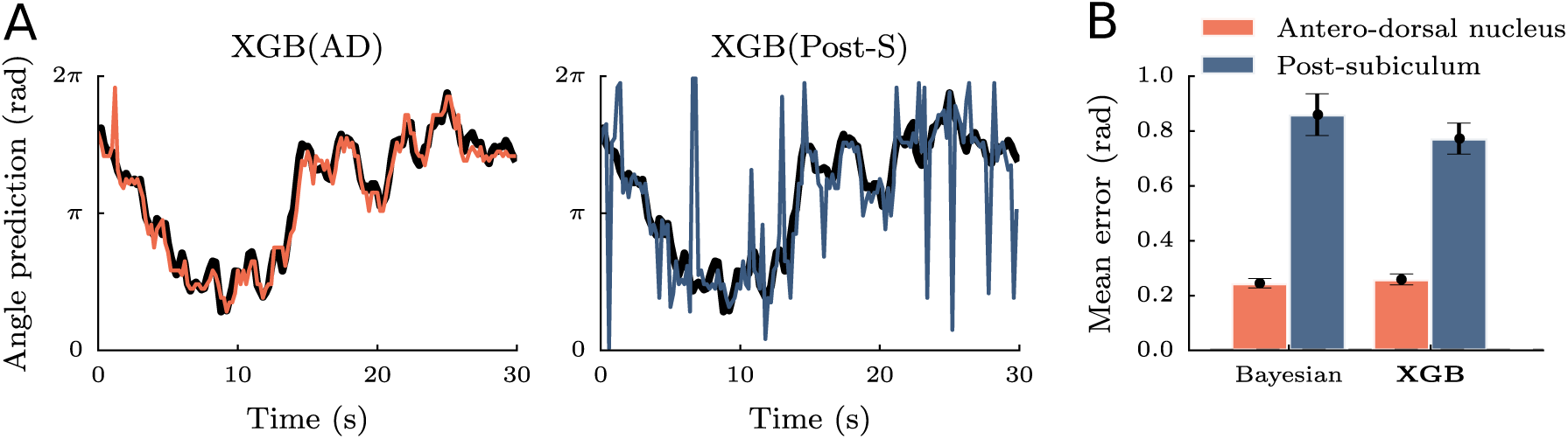
Decoding of HD angle. **A** Example of decoding for XGB during 30 seconds of head rotation for both ADn and PoSub spiking activities. The black line shows the real angular HD.**B** For sessions with large groups of neurons (n ≥ 7) in ADn and PoSub, the HD of the animal was decoded based on spiking activity with the classical Bayesian decoding and gradient boosted trees (XGB) over 60 angular bins.

**Figure S2:**
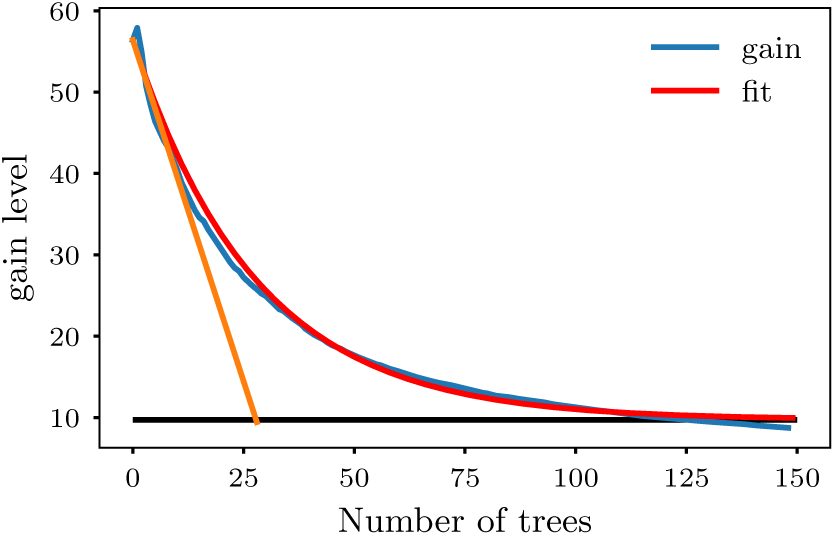
Learning gain per tree decreases with the number of trees (blue line). This decay was well captured by an exponential fit (red line), from which an optimal number of trees of approximatively 30 trees is derived (intersect of the linear fit at origin with the x-axis). At this stage the mean gain per tree is approximately 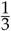 of its initial value and most of the learning has already occurred.

**Figure S3:**
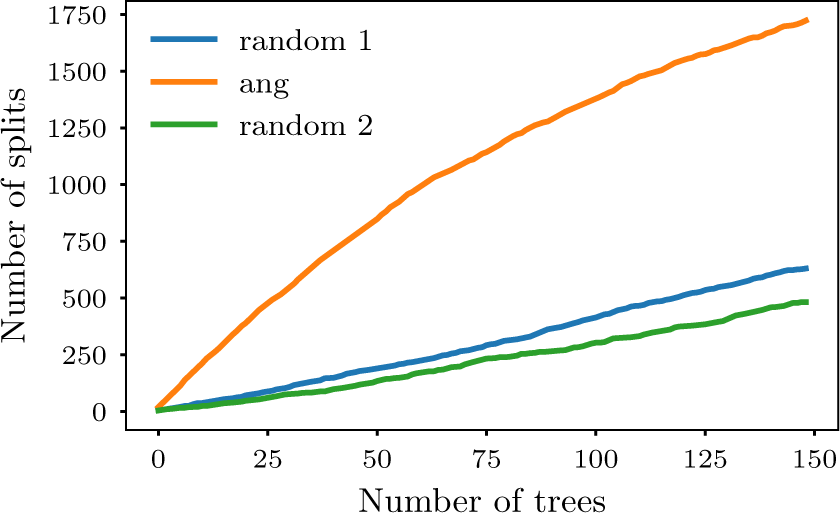
Features carrying actual signal are preferentially split in the first trees, resulting in higher gain. The graph illustrates the evolution of split density when learning the spike train of a HD neuron as a function of the number of trees for three features: the actual HD and two random vectors. Split density increased linearly and similarly with the number of trees in the asymptotic regime for all features. However, the increase was much higher for the at low tree numbers, a difference well captured by gain analysis. Note that, as the order of features in the algorithm may impact which are split first, we showed how the feature data were organized (random 1, angle and random 2).

**Figure S4:**
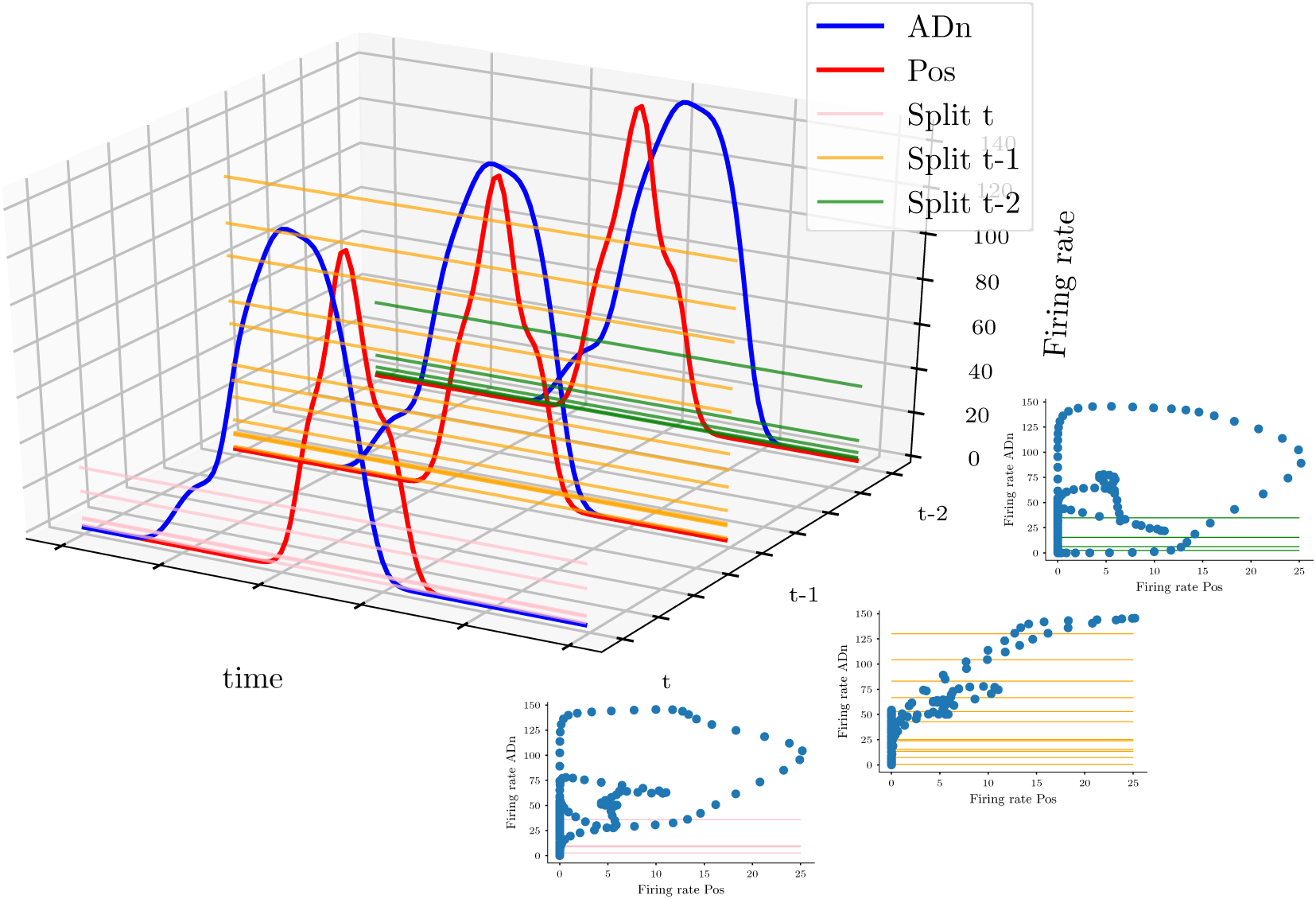
Revealing temporal delay in peer-prediction. Feature space is composed of multiple copies of the activity of the feature neuron (in this case, in the ADn) at various time-lags (blue curves) to learn the target spike train (PoSub, red curves). The relationship between the two spike trains shows maximal dependence at t-1, resulting in a high number of splits by the algorithm (yellow horizontal lines). Splitting was less effective for more independent firing at t and t-2. In this example, the relationship at t-1 is trivial (linear and positively correlated). However, the quantification of these interactions give comparable values for a large variety of interactions (e.g. positive, negative or monotonically non linear).

